# Nucleotide-level chemical reaction network modeling enables quantitative prediction of reconstituted cell-free expression system

**DOI:** 10.64898/2026.02.22.707325

**Authors:** Zoila Jurado, Ayush Pandey, Richard M. Murray

## Abstract

Cell-free expression systems offer a method for rapid prototyping of DNA circuits and functional protein synthesis. While crude extracts remain a black box with many components carrying out unknown reactions, PURE contains only the required transcription and translation components for protein production. All proteins and small molecules are at known concentrations, enabling detailed modeling for reliable computational predictions. However, there is little to no experimental data supporting the expression of target proteins for PURE-based models. In this work, we generalized the PURE detailed translation model for proteins with arbitrary amino acid compositions and lengths. We then built a chemical reaction network for transcription in PURE, validating the transcription models using DNA expression for the malachite-green aptamer (MGapt) to measure mRNA production. Lastly, we coupled the transcription and the generalized translation models to create a PURE protein synthesis model built purely of mass-action reactions. We used the combined model to capture the kinetics of MGapt and deGFP expressed from plasmids at varying concentrations.

## Introduction

Cell-free protein synthesis (CFPS) systems are divided into two broad categories. The most commonly used CFPS is based on cell lysate, first implemented in the 1960s to express synthetic RNAs to decipher the genetic code [1]. Cell lysate-based systems or TX-TL (transcription and translation) utilize cellular machinery harvested from the cell. After multiple stages of growth, lysis, and clarifying spins, an energy buffer with amino acids is added to the lysate to create a protein expression system [2, 3]. The widespread use of TX-TL, now commercially available, has its limitations. The research field is limited by batch-to-batch variability, affecting lifetime and total protein expression [4]. The batch-to-batch variability can result from multiple variables, such as cell strain, optical density (OD) at the time of harvest, lysis method, energy mixture composition, and reagent batches. The ability to achieve ‘design–build–test’ cycles, similar to those found in other traditional engineering fields, is thus ultimately limited by the variability of the CFPS system.

One advantage of using cellular lysate is the retention of biological pathways of the cell strain, such as glycolysis, facilitating energy regeneration. Unfortunately, the extent to which cellular processes remain functional is still undetermined. Over the last decade, multiple TX-TL protein expression models have been proposed and have been shown to capture RNA and protein production [5, 6, 7, 8]. However, these models cannot predict expression without characterizing them to their specific experimental data sets. Modeling the behavior of whole circuits using a software toolbox and characterizing components of the entire model [9] are steps towards more predictive models. While these TX-TL models can help understand phenomena or estimate unknown parameters, they are constrained by the unknown composition of the cellular lysate. Characterization of lysate as the next step for TX-TL modeling relies on LCMS to measure small molecules, proteins, and lipids to understand how much of the core metabolism is active, potential side reactions, and waste generation effects [10, 11]. However, measuring all proteins, small molecules, and lipids, then mapping the chemical reactions associated with each lysate, is currently not feasible due to the lack of lysate standardization. Thus, having a universal, batch-independent, and detailed TX-TL model is far off.

In contrast to cellular lysate-based cell-free protein synthesis, purified components can be used to express functional proteins. The first reported attempt was in 1977 [12] with limited success. Weissbach’s group provided a starting point to which Ganoza *et al*. [13], and Pavlov *et al*. [14, 15] attempted to use precharged aminoacyl-tRNAs or partially purified aminoacyl-tRNA synthetase alongside purified proteins. Shortly after, in 2001, Shimizu *et al*. [16] achieved successful protein production using PURE — Protein synthesis Using purified Recombinant Elements. To recreate the “central dogma,” PURE contains all transcription and translation proteins required for protein production at known concentrations. Consequently, enabling batch-to-batch and inter-laboratory repeatability and the opportunity for detailed modeling for reliable computational predictions. Nonetheless, even with complete control and knowledge of the composition, existing PURE models fall back to the phenomenological modeling of transcription and translation [5, 17, 18, 19, 20] by grouping all NTPs as one variable, not modeling each step of protein production, and employing Hill functions instead of chemical reaction equations. Although such phenomenological models are simple and capture overall expression levels, they are only valid under specific assumptions and not readily extensible to model experimental conditions in which specific components are varied.

In 2017, Shimizu’s group introduced a MATLAB model for the translation mechanisms in PURE [21, 22], which will be referred to as the ePURE-MATLAB model. The proposed detailed model distinguishes between NTPs and includes each elongation step in the growing peptide. This computational model comprises 968 mass-action reactions and 241 species, of which 27 components initialize the PURE model. Time courses of all components can be tracked in this model, which provides a valuable method to explore and systematically model protein synthesis in PURE. Even so, the model only simulates the expression of a small peptide, fMGG, with each reaction explicitly written on a spreadsheet, and cannot be easily extended to longer proteins. In 2023, Shimizu’s group introduced ePURE_JSBM, which extended the framework to model peptide elongation from mRNA [23]. However, because the ePURE-JSBM model only includes peptide elongation and omits transcription, it provides an incomplete description of PURE that cannot be directly validated against experimental measurements of coupled transcription-translation. In this paper, to address these limitations, we demonstrate (1) a translation model for arbitrary proteins; (2) a detailed model of PURE for the transcription of arbitrary DNA sequences with mechanistic details for each step in the transcription parameterized with malachite-green aptamer (MGapt) expression in PURE; and a coupled transcription and translation model of PURE that ultimately captures kinetic trends of MGapt and a green florescence protein (deGFP). By quantitatively capturing both transcription and translation kinetics from first principles, this model reveals how molecular-level mechanisms propagate to determine system-level expression outcomes in reconstituted biochemical systems.

## Results and Discussion

### Generalization of a PURE Translation Model for Proteins of Arbitrary Amino Acid Composition

The ePURE-MATLAB PURE translation model [21, 22] is limited to the expression of the fMGG peptide. The model comprises 968 reactions and 241 species, all explicitly written. As a result, the ability to change or extend the peptide is laborious and futile. To make peptide variation more tractable, we used a CRN compiler tool called BioCRNpyler [24]. BioCRNpyler is a Python-based software package that can compile CRN models from a high-level description of a system. BioCRNpyler provides an object-oriented, modular framework to compile components and the mechanisms that describe the interactions between the components into a CRN model by automatically generating all reactions, species, and parameters. The CRN model can be exported as SBML, and the constituent components and associated parameters can be shared to build other larger system models. Thus, building the PURE model with BioCRNpyler enables a framework that can be easily modified, expanded, and shared in other models. BioCRNpyler also contains a library of parts and parameters that can be used to share parts of the model in other larger system models. We first convert the ePURE-MATLAB translation model to Python using BioCRNpyler for the peptide fMGG. Using the spreadsheet-based ePURE-MATLAB model, the reactions are split into four groups: initiation, elongation, termination, and auxiliary reactions. To generate species and reactions dependent on the desired peptide, we leverage features in BioCRNpyler that enable scalability essential for the elongation step. Retaining all the reactions and parameters from the ePURE-MATLAB model, the PURE-TL model loops to iterate over peptides of arbitrary length. The comparison between the PURE-TL model (magenta line) and ePURE-MATLAB model (blue circles) can be seen in Figure 1a, with the error between the two in orange. The absolute error remains below 0.2 % throughout the time course, demonstrating that the BioCRNpyler framework accurately reproduces the ePURE-MATLAB predictions.

**Figure 1:**
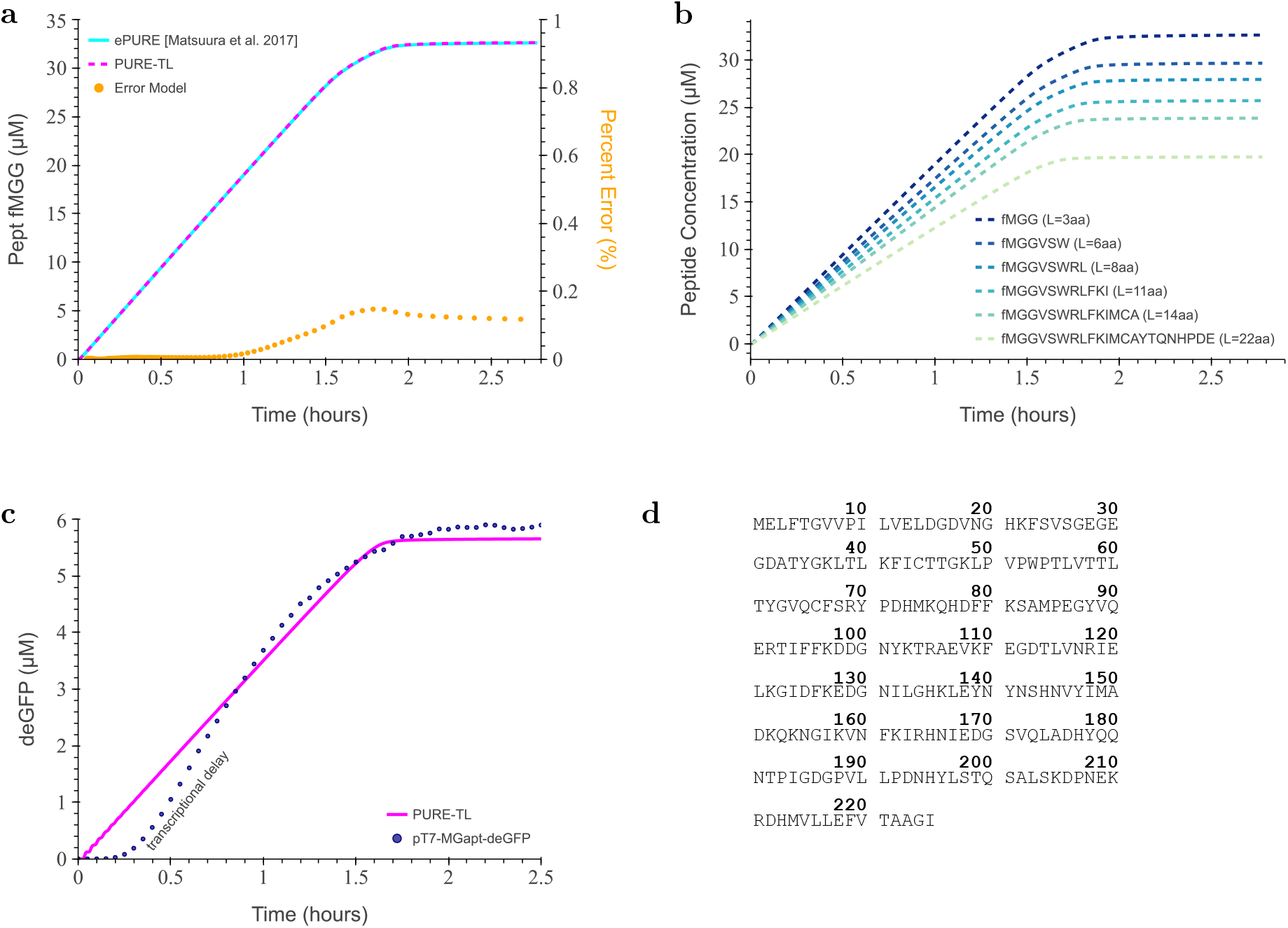
Generalization of the PURE-TL model and comparison against expressed deGFP from mRNA. (**a**) Comparison of original ePURE-MATLAB model to PURE-TL model. PURE-TL model (magenta line) and ePURE-MATLAB model (blue circles) overlap, and the difference between models (orange circles) is shown on a secondary axis. (**b**) Generalization of the PURE-TL model with different amino acids to the original fMGG peptide. (**c**) Simulated translation of 0.126 µm RNA expressing deGFP (magenta line) overlaying experimental production of deGFP from DNA at 5 nm (blue circles). (**d**) Amino acid list used as input for the PURE-TL model.

The generalized PURE translation model in BioCRNpyler (PURE-TL) was built by adding different amino acids one by one until all 20 amino acids were incorporated (see Figure 1b). As expected, the amount of the final peptide decreases with increasing amino acid chain length as a consequence of resource constraints. Next, the model was expanded for repeated amino acids and arbitrary amino acid sequences of length greater than 21. Finally, the translation of deGFP (RNA_*n*_=805, Pept_*n*_=226) was modeled using the initial conditions adopted from the ePURE-MATLAB model. We set the initial conditions for tRNAs and amino acids not associated with Met or Gly equal to the initial conditions of the Gly-amino acid and tRNA as previously given in Table S4. We initialed the PURE-TL with RNA_*n*_ to 0.126 µm, such that the total deGFP expression is comparable to experimental results of 6 µm, as seen in Figure 1c. The discrepancy within the first hour of protein expression, indicated by the transcriptional delay label, is expected because the simulation is initiated with a nonzero amount of RNA, causing deGFP production to begin immediately. Experimentally, however, RNA must first be transcribed from DNA before translation can occur. Modeling the translation of an experimentally relevant protein, such as deGFP, therefore enables direct comparison between the PURE model and experimental results.

#### Validation of translation model

To verify the extended translation model, we measured deGFP production of 10 µL PURE reactions with purified RNA of MGapt-UTR1-deGFP at final concentrations of between 0.22 µm and 3.38 µm (see Supplementary Information Section 2.2). To capture the non-linear relationship between RNA and protein production, observed in Figure S3, we propose an RNA effective multiplication factor, RNA_effective_. The RNA_effective_ factor serves as a phenomenological correction to account for processes that reduce translation efficiency but are not explicitly represented in our mechanistic model, such as waste product accumulation.

Utilizing the RNA effective multiplication factor, we modeled the translation of deGFP using purified RNA at 0.41 µm, 0.86 µm, 1.26 µm, 1.67, and 2.53 µm. Figure 2 shows the comparison between the simulated translation deGFP production model (solid lines) and the experimental data (circles with error bars) at various start RNA concentrations in corresponding colors.

**Figure 2:**
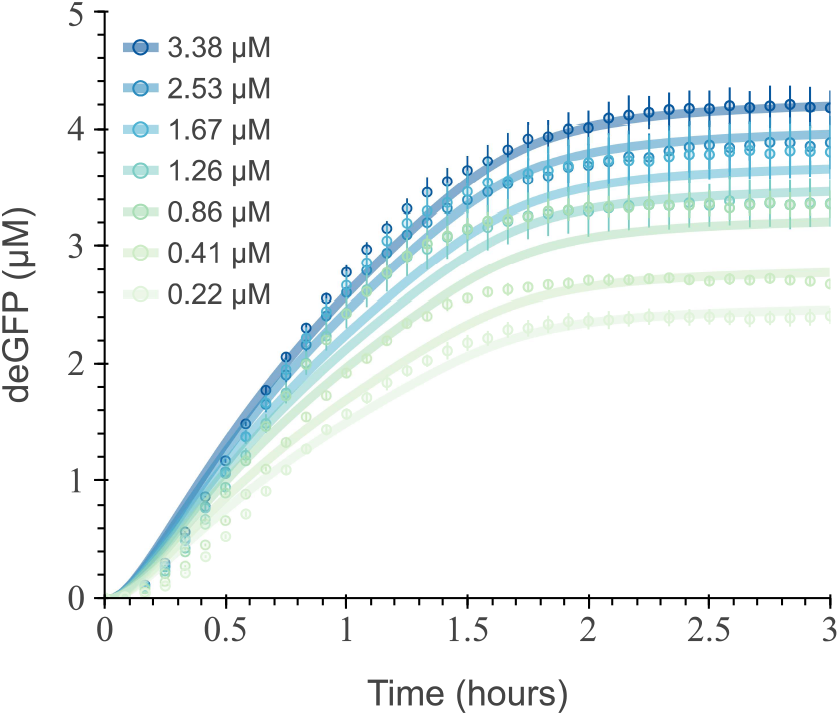
Simulated deGFP production with and without leveraging effective RNA calculation. Simulated deGFP production of PURE-TL model initialized at various RNA concentrations overlaying experimental results (circles with error bars) at respective color concentrations (solid lines), using effective RNA concentration as described in Supplementary Information Section 3.10.

The translation model captures the final deGFP production with an average error less than 3.74 % (see Table S1). The model overestimates the deGFP production rate, with the simulation predicting faster protein accumulation than is observed experimentally. The disparity between the translation model and experimental data suggests that tuning the translation model parameters would be necessary. Due to the extensive size and time required to fit over 1000 reactions, we opted to continue using the published parameters. Future work will focus on further parameterization of the translation parameters.

### A Chemical Reaction Network for PURE Transcription

Having established a translation model for arbitrary protein sequences, we next developed a corresponding transcription model to capture mRNA synthesis. To model the detailed step-by-step mechanistic transcription process, we built a chemical reaction network (CRN). We developed a generalized BioCRNpyler framework that automatically constructs reaction networks for arbitrary DNA sequences based on the mechanistic transcription mechanisms characterized by Tuza *et al*. [25]. This tool-based approach requires users to input the coding sequence for their desired protein (excluding promoter and terminator regions). Then the complete set of species and reactions needed to simulate transcription dynamics is automatically generated.

We split transcription, using the translation model as a framework, into three groups: initiation, elongation, and termination, as illustrated in Figure 3 for RNA of length n. Initiation steps include a one-step GTP-dependent activation of T7 RNAP [26]:

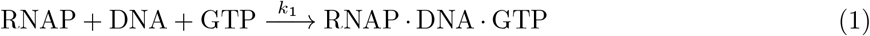

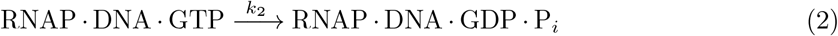

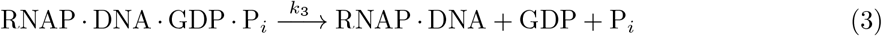

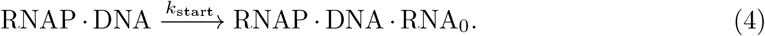

**Figure 3:**
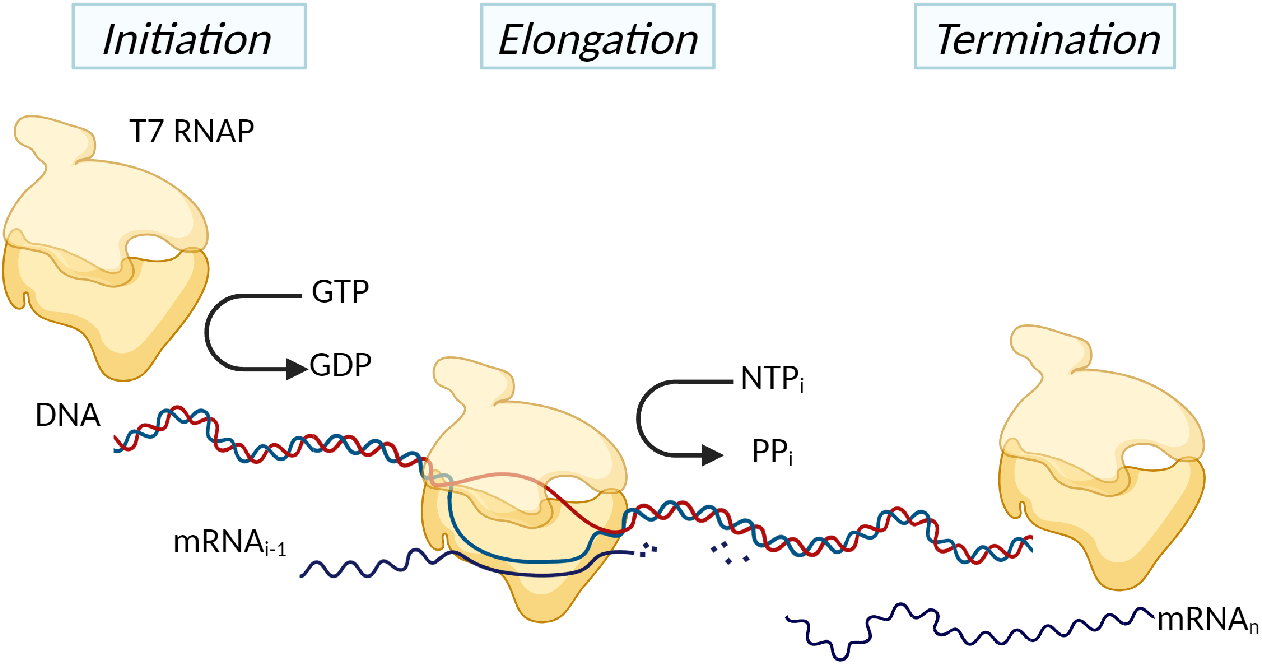
Schematic of RNA synthesis with a reconstituted *E. coli* transcription system using T7 RNAP. We split the transcription reactions into three sub-processes: initiation, elongation, and termination. Auxiliary reactions, such as energy recycling or those explicitly related to translation, are not included. Created with BioRender.com.

Elongation steps model each binding state separately for the addition of an NTP:

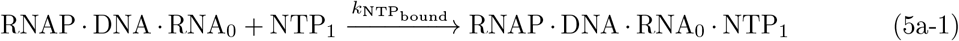

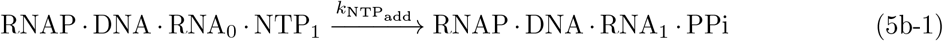

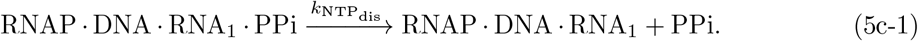

The elongation steps repeat for each nucleic acid addition along the growing RNA chain of length *n*:

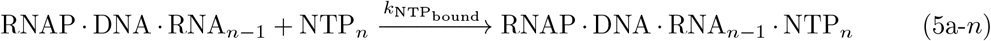

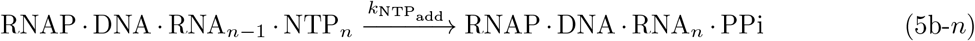

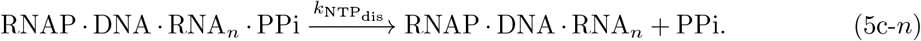

Finally, the termination step models the dissociation of the RNA_*n*_, DNA, and T7 RNAP:

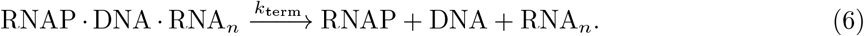

In our transcription reactions, we do not explicitly account for multiple polymerases simultaneously bound to a single DNA strand. However, the effects of simultaneous transcription are captured in the reaction rates. Additionally, no auxiliary reactions, such as NTP recycling, are incorporated into the transcription model.

#### Parameter inference on the transcription model

We measured the T7 RNAP-driven transcription of MGapt without a ribosome binding site (RBS). The lack of RBS limits the reactions associated with translation, allowing the focus to be on transcription. PURE reactions at 10 µL were conducted using a plasmid DNA construct of pT7-MGapt-tT7. The total amount of RNA was calculated using dynamic calibration curves in Figure S2 for the respective construct.

Before starting the parameter inference pipeline (see Materials and Methods), we account for translation reactions independent of peptide synthesis, such as tRNA charging. We modify the initial conditions for the transcription model of DNA construct pT7-MGapt-tT7 in the Bayesian inference pipeline. To determine the amount of ATP and GTP consumed by these reactions within the first 15 min, the translation model was run without DNA. The initial conditions were taken from PURE components [27] for NTPs and T7 RNAP concentrations. The initial conditions for the transcription-only model are given in Table S2.

To parameterize our transcription model, we initialize the parameters using the values from Tuza *et al*. [25]. Training our model using parameter identification is not straightforward, since we only have experimental data for MGapt fluorescence. Therefore, it is crucial to assess the empirical identifiability of the model parameters. To assess identifiability and determine posterior parameter distributions, we utilize a biological data analysis pipeline [28] implemented in the Python package BioSCRAPE [29]. This pipeline provides a practical interface to the BioCRNpyler model to run these analyses. Next, to account for the intrinsic noise observed in the experimental data, we use Bayesian inference to obtain a distribution of possible parameter values given the experimental data.

We choose to fit all eight reaction rates in equations (1)-(6) using a coarse tuning to search within the parameter space equal to twice the initial parameter given in Table S3, based on Figure 4a. The initial posterior distributions of parameters, shown in Figure S5, were obtained using the Bayesian inference tools in BioSCRAPE on all the reaction rates. Subsequently, the model was retrained using a narrower standard deviation around the results from the initial inference. The corner plot in Figure 4b shows the posterior parameter distributions and their covariance. This chart provides a distribution to sample from when predicting the output using the fitted model. The model simulation with parameter values drawn from the posterior is shown in Figure 4c, along with the experimental data.

**Figure 4:**
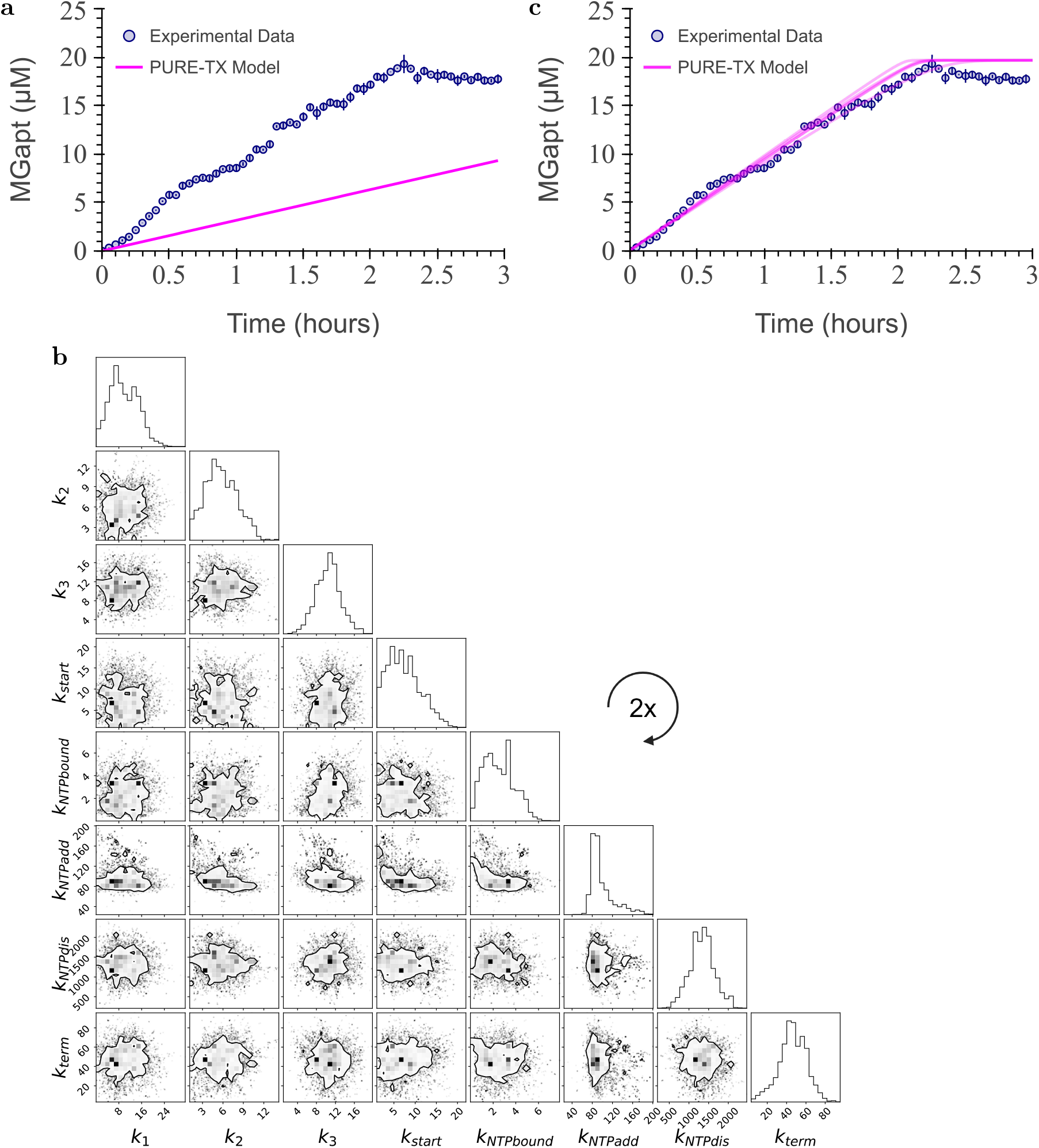
Modeling, analysis, and parameter learning for the PURE transcription-only model. **(a)** The model simulations (in magenta) with the original reaction parameters are compared with the experimental data for three biological replicates (blue circles with error bars). (**b**) The posterior distributions of parameters were obtained after running Bayesian inference on all eight reaction rates. The corner plot depicts the covariance of the eight parameters, with the contour showing the 75 % probability region for the parameter values. (**c**) Model simulations (in magenta) with parameter values for all reaction rates sampled from the posterior distributions are shown alongside the experimental data for three biological replicates (blue circles with error bars).

Transcription does not occur independently; the presence of translational components influences transcription, even when mRNA-specific translation is absent. To further validate our transcription model, we proceeded to a coupled transcription-translation system. We reintroduce the auxiliary translation reactions associated with energy regeneration, tRNA charging, and ribosome interactions. Next, we ran local sensitivity analysis for a expression of MGapt from plasmid pT7-MGapt-tT7 of all species in the PURE transcription-only model against all parameters. The sensitivity analysis heatmap is shown in Figure 5a, from which we select the most sensitive parameters for MGapt production. Out of the eight reaction rates highlighted in equations (1)-(6), we find that the most sensitive parameters are *k*_2_, *k*_3_, *k*_start_, and 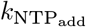. These parameters correspond to the initiation of transcription and consecutive addition of NTPs to the growing RNA strand, respectively. The initial conditions for the transcription and translation model are given in Table S4; based on Version 7 PURE concentration published in Table S1 by Kazuta *et al*. [30].

**Figure 5:**
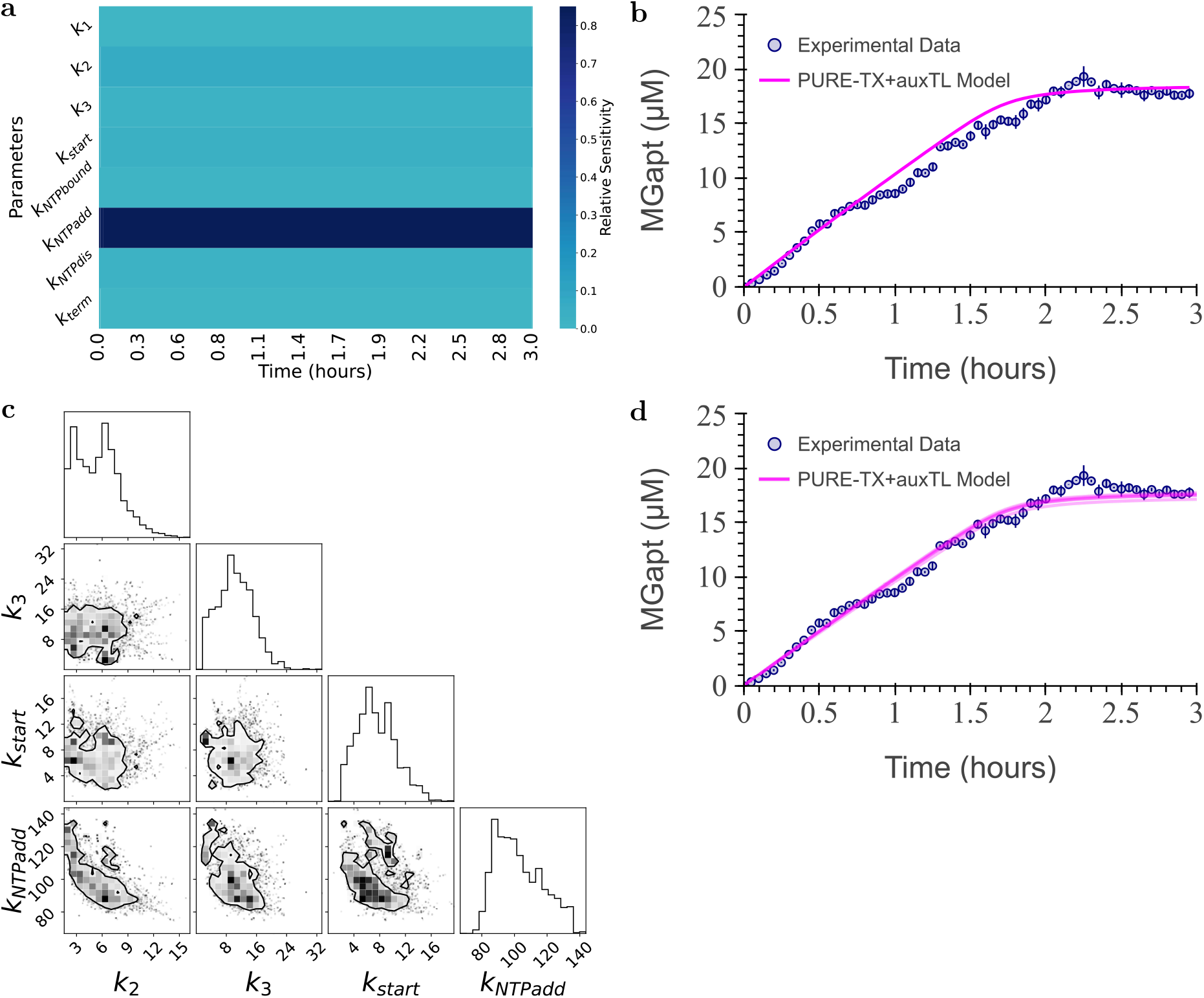
Modeling, analysis, and parameter learning for the PURE model without protein production. (**a**) The sensitivity of the MGapt fluorescence to the PURE transcription-only model parameters at all times. (**b**) The PURE model simulations (in magenta) with the previously fitted reaction parameters and the experimental data for three biological replicates (blue circles with error bars). (**c**) Based on sensitivity analysis in (a), we use the MGapt fluorescence to infer the posterior parameter distributions for *k*_2_, *k*_3_, *k*_start_, and 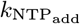. The corner plot depicts the covariance of the four parameters, with the contour showing the 75 % probability region for the parameter values. (**d**) Parameter values drawn from the posterior distributions shown in (c), the model predictions (magenta) are shown alongside the experimental data. Parameter values for all reaction rates sampled from the posterior distributions, the model simulations (in magenta) are shown alongside the experimental data for three biological replicates (blue circles with error bars). The simulations includes auxiliary reactions related to translation components but without mRNA specific translation.

Following a similar pipeline as before, we use Bayesian inference to identify the posterior parameter distributions for these four parameters. The PURE simulation of the MGapt production using the reaction parameters from the translation model is shown in Figure 5b. The corner plot in Figure 5c shows the posterior parameter distributions and their covariance. The model simulations using parameter values drawn from the posterior and experimental data are shown in Figure 5d. The final parameter values used are given in Table S5.

Summarizing the computational analysis, we utilized transcription parameters based on results from the literature to fit a transcription-only model against MGapt expression in PURE. Then, we inferred the most sensitive parameters *k*_2_, *k*_3_, *k*_start_, and 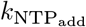 in the transcription model containing auxiliary translation reaction. The final trained model accurately predicts the transcription activity of the DNA sequence for which the transcription reactions were generated. Notably, the coupled TX+auxTL model (Figure 5) achieves better quantitative agreement with experimental MGapt concentrations than the transcription-only model (Figure 4). This improvement demonstrates that accurate modeling of transcription kinetics requires accounting for the auxiliary translation reactions that both regenerate and consume energy substrates.

### PURE Model: Coupled Transcription and Translation Models

We use BioCRNpyler to combine the separate transcription and translation models for arbitrary sequences. Reactions and species from the translation and transcription models are compiled together, and any duplicate species, such as ATP, GTP, and PPi, are removed. Next, the transcription model’s output, RNA_*n*_, and translation model’s input, RNA, were linked with the uni-direction mass-action reaction:

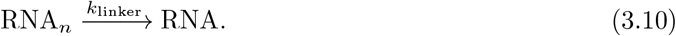

The parameter value of *k*_linker_ is set to 1000 s^−1^, such that all of the RNA produced in the transcription model will be instantly available for translation. In this coupled framework, the total RNA output from transcription serves as input to translation, eliminating the need for the phenomenological RNA_effective_ factor required in the standalone translation model. The inclusion of the reaction in equation (3.10) can be omitted by utilizing the same species as the output and input of the respective models. However, this linker reaction provides a modular framework for future incorporation of RNA degradation, diffusion limitations, or other transport processes that may affect RNA availability for translation.

To account for protein folding, the following reaction was added:

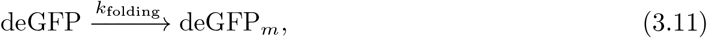

where *k*_folding_ estimated as 600 ^−1^ s^−1^ [31], completing the combined PURE model. The total number of reactions of the combined transcription and translation model is 6988 with 6280 species. Unable to obtain the initial concentrations of NEB PURExpress, the initial concentrations were set to the Version 7 PURE concentration published in Table S1 [30]. The one exception is that the small molecule creatine phosphate (CP) has an initial concentration of 10 mm; see Table S4 for the initial conditions of all of the proteins and amino acids.

In Figure 6a, RNA production is consistent with experiments until approximately 1 h. After 1 h the model over predicts MGapt production by approximately 40 %. The difference after 1 h in total MGapt produced between the predicted and experimental data may be attributed to Mgapt degradation or inaccuracies in the initial conditions, such as the model having more total available energy than is actually present or usable in the experimental system. Additionally, the slight delay observed in the simulated MGapt and experimental MGapt is likely due to the time elapsed between mixing the reaction and transferring the plate to the plate reader.

**Figure 6:**
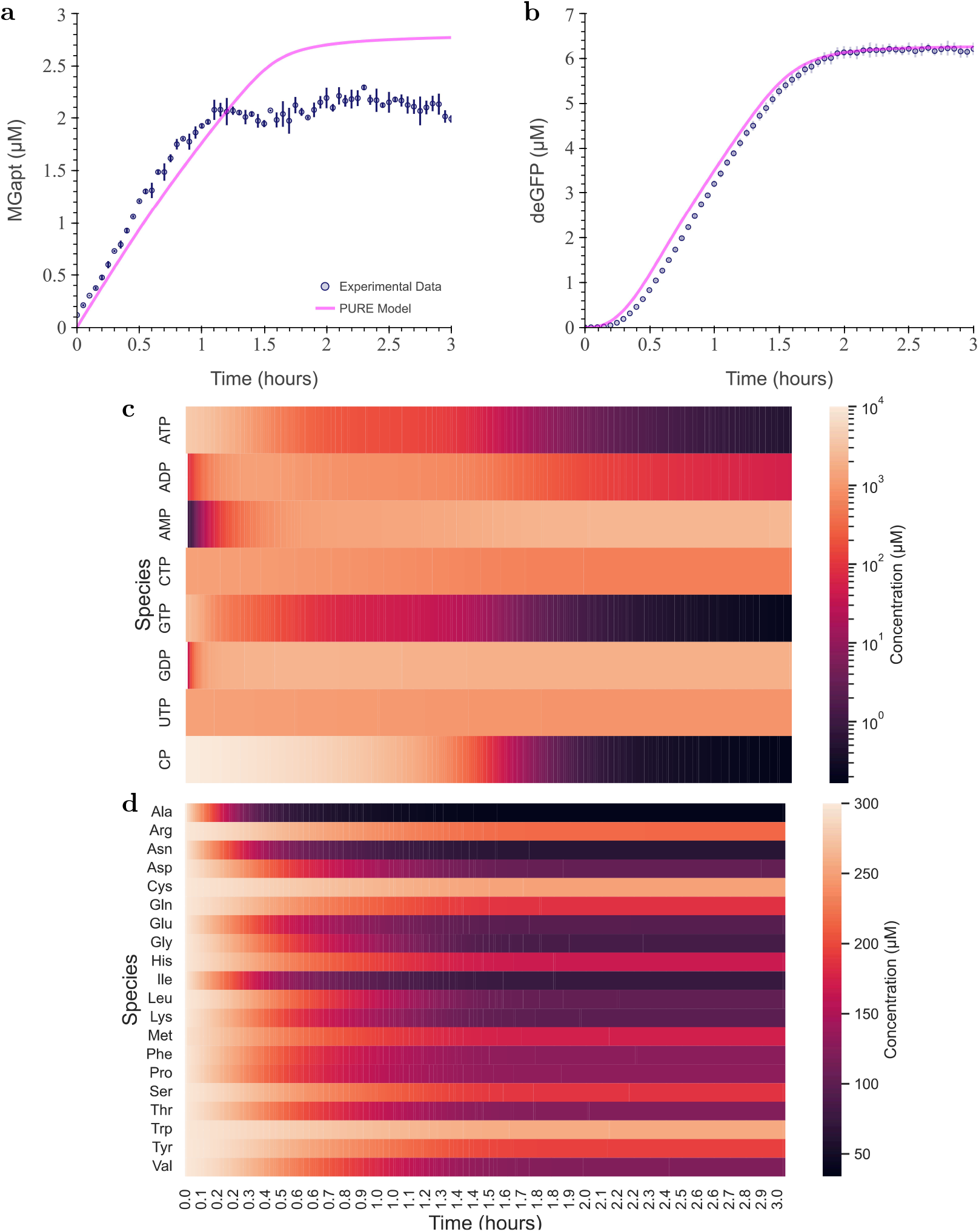
The combined transcription and translation model for pT7-MGapt-UTR1-deGFP-tT7, DNA=5 nm, with experimental result in PURExpress. (**a**) Modeled RNA production in the combined model (magenta line) overlaying with experimental data, three replicates (blue circles and blue error bars). (**b**) Modeled deGFP expression in the combined model (magenta line) overlaying experimental data, three replicates (blue circles and blue error bars). (**c**) Concentrations (µm) of ATP, ADP, AMP, CTP, GTP, GDP, and UTP over simulation time using log-normal scale. (**d**) Concentrations (µm) of all amino acids over simulation time. Simulations use a protein folding efficiency of 0.7.

The production of deGFP from DNA construct pT7-MGapt-UTR1-deGFP-tT7 at 5 nm did not initially capture deGFP production despite the model’s accuracy of RNA produced within the first 1 h. The over-prediction of deGFP may be the lingering non-linear correlation between RNA and deGFP production observed earlier. However, any inhibition of overloading the translation reaction with RNA would not be as significant, as RNA is being slowly produced. A more plausible alternative explanation is that incomplete protein translation occurs in PURE cell-free protein synthesis systems (see Supplementary Information Section 4.1), leading to diminished functional protein compared to what is theoretically possible. To account for misfolding, premature termination of translation, and aggregation, we reduce the simulated deGFP production by 30 %.

The expression pattern of deGFP in the coupled transcription-translation CRN model more closely resembles actual experimental data in terms of protein expression dynamics, compared to the translation-only model shown in Figure 1c. In Figure 6b, deGFP production stops with a smoother transition compared to Figure 1c. By including the scaling factor of 0.70, the PURE model accurately predicts total deGFP production throughout the reaction’s lifetime with a 5 % error, as shown in Figure S7. The consistency of this factor across DNA concentrations suggests it represents a fundamental limitation of translation efficiency in PURE rather than a condition-dependent correction. Moreover, since the model captures early-time mRNA dynamics accurately (approximately 1 h), simply adjusting transcription parameters to better fit mRNA kinetics would not resolve the deGFP overprediction without the correction factor.

Finally, due to the detailed nature of the model, we can track the concentrations of all proteins, amino acids, and energy carriers and begin exploring the limiting factors of PURE. For example, we can use heatmaps of the concentrations of energy-related small molecules over time, in Figure 6c, to hypothesize that ATP, GTP, and CP are the likely limiting energy carriers in PURE. Specifically, the consumption of GTP and ATP likely limits RNA and protein production. We can also speculate that the energy source, CP, which serves as a phosphate donor for ATP synthesis, is fully depleted by 2 h. Additionally, Figure 6d shows that amino acids are not limiting in PURE. The excess of amino acids is apparent, with none of the 20 amino acid concentrations falling below 20 µm.

#### Validation of combined transcription and translation model

To validate the PURE model, which combines the transcription and translation model, we ran additional 10 µL PURE reactions with DNA of pT7-MGapt-UTR1-deGFP-tT7 at final concentrations of 0.07 nm, 0.12 nm, 0.26 nm, 0.47 nm, 1.06 nm, and 1.99 nm. We observed that the relationship between initial DNA concentration and deGFP production at 2 h was non-linear (see Figure S8a). Specifically, we note diminishing protein production as DNA concentration exceeds 0.5 nm. To further investigate the origin of the nonlinearity, we plotted the relationship between DNA added to MGapt produced and Mgapt produced to deGFP produced in Figure S8b and Figure S8c, respectively.

The relationship between initial DNA concentration and MGapt produced in Figure S8b is linear at DNA concentration above 0.5 nm, indicating a change of transcriptional regime at higher DNA concentrations. The shift in the transcription regime may indicate a transition from a DNA-limited regime. However, Figure S8c shows a diminishing return on protein production when the maximum concentration of RNA produced surpasses 1 µm. The diminishing return of deGFP relative to RNA, previously observed in the translation experiments, may result from the saturation of translation proteins, production of inhibitory products, or reduced levels of ATP, GTP, and other small molecules not currently incorporated in the model.

Notably, in the translation results shown in Figure S3, when purified RNA was added at 0.855 µm and 2.528 µm, the respective deGFP expression was 3.08 µm and 3.78 µm. This is approximately 60 % lower than the deGFP expressed at comparable synthesized RNA concentrations in Figure S8c. These results suggest that starting with DNA in the PURE reaction is more efficient for protein production than dosing in purified RNA. While the precise mechanism requires further investigation, we hypothesize that increased RNA concentrations lead to competition for limited translation machinery and energy substrates, reducing the total amount of proteins fully expressed. Consequently, similar to the validation of the translation model, we propose using an effective DNA (DNA_effective_) to recapitulate the maximum RNA synthesized experimentally in the model (see Supplementary Information Section 4.4).

Using a DNA effective multiplication factor, we modeled the transcription and translation of deGFP at the different DNA concentrations. The simulation results are shown in Figure 7 and Table S6. In Figure 7, the simulation results (solid lines) are superimposed on experimental data (circles and error bars) with the varying DNA concentrations represented by different shades of purple in descending order.

**Figure 7:**
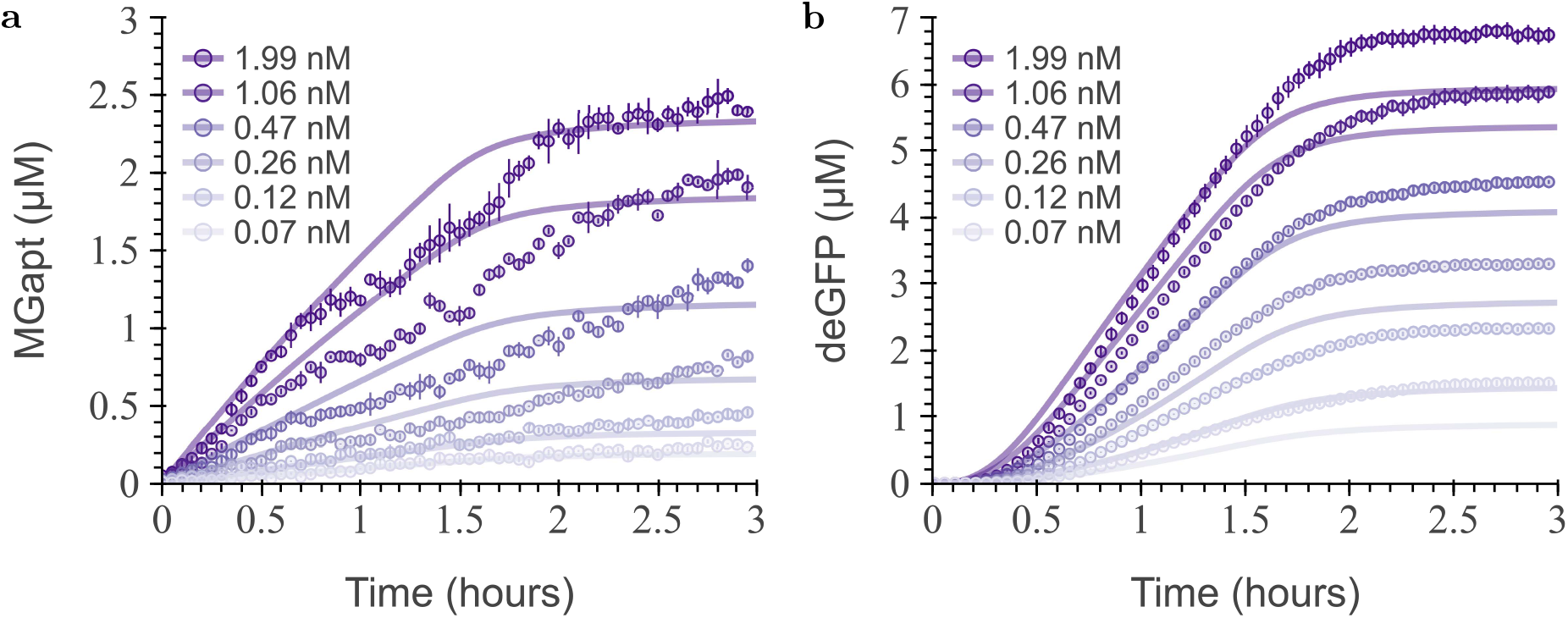
The PURE model for pT7-MGapt-UTR1-deGFP-tT7 leveraging effective RNA calculation, at different initial DNA concentrations compared with experimental results in PURExpress. Simulations use effective DNA concentration as described in Supplementary Information Section 4.4 and a protein folding efficiency of 0.7. The simulation results (solid lines) are overlayed with experimental data (circles and error bars) for the production of (**a**) RNA and (**b**) deGFP. The different DNA concentrations are reflected in the different shades of purple in decreasing order.

Although the simulated result does not capture the MGapt non-monotonic dynamics depicted in Figure 7a, the model approximately agrees with the experimental results within the first 45 min and the total MGapt expressed. In Figure 7b, the model typically underestimates the final deGFP concentrations, but mirrors deGFP dynamics up to 1.5 h at DNA concentration above 0.47 nm. Further fitting of the translation parameters would be preferable; yet, we remain constrained by the unknown compositions of PURE. Uncertainty in the concentrations of essential proteins and contaminants may result in unaccounted-for reactions. However, despite reactions that still need to be identified and incorporated, the PURE model remains useful for understanding protein production in PURE systems. The model can capture general trends and approximate the coupled expression of MGapt and deGFP, thereby providing insight to help guide the design of experiments to test hypotheses.

## Conclusion

We have proposed a coupled transcription and translation model of cell-free protein expression for arbitrary proteins in PURE. The existing model of translation in PURE from Matsuura *et al*. [21] only modeled the translation of the fMGG peptide. In this paper, we have generalized a PURE translation model to any arbitrary protein. Next, we developed a transcription model that incorporates each step of the growing mRNA strand and the specific NTP required to transcribe the mRNA from any given DNA sequence. Finally, we combined the transcription and translation models to obtain a holistic model of protein expression in PURE. Using our approach, it is possible to create mathematical models of the expression of experimentally relevant proteins in PURE.

We validated our models using experimental data. We identified a distribution of possible parameters in the model with the experimental data for MGapt fluorescence. We demonstrate that the validated model accurately predicts MGapt fluorescence for two different plasmids — one without an RBS and one co-expressing deGFP. The combined model of the PURE cell-free system, using the validated transcription model and the extended translation model, was then used to predict deGFP expression. The model predictions are shown to approximately agree with the experimental results at DNA concentrations around 0.5 nm and higher.

While our model captures detailed mechanistic steps in transcription and translation, accurate experimental prediction required phenomenological scaling factors (RNA_effective_ and DNA_effective_) rather than relying solely on first-principles mechanisms. These corrections highlight the incompletely characterized loading effects on transcription and translation machinery, including resource competition, saturation dynamics, and inhibitory product accumulation. Future improvements should incorporate explicit reactions for inhibitory species (inorganic phosphate, pyrophosphate, pH effects), DNA replication, ribosomal loading dynamics, and incomplete transcription/translation products [32]. Such mechanistic refinements could obviate the need for empirical scaling factors and achieve a fully predictive, first-principles model of cell-free gene expression.

This detailed model of the PUREexpress system is a step towards cell-free protein synthesis CFPS characterization. The model provides a platform for testing multiple hypotheses to guide experimental design in PURE. Further development of the model with OnePot PURE [33], a version of PURE where all 36 proteins are co-cultured and purified together, can help circumvent batch-to-batch and inter-laboratory variability problems seen in extract-based systems. Beyond this, the model can be instrumental in identifying potential directions for further research, particularly regarding the coupling between transcription and translation mechanisms. We hope this model will bring together multiple research groups and facilitate the exchange of information to support characterization and continued optimization. Together, we can build a library of characterized parts for PURE or OnePot PURE to achieve a robust ‘design–build–test’ cycle.

## Materials and Methods

### Computational Modeling and Simulations

In this paper, we build on a previously published deterministic model of protein synthesis of a formyl-Met-Gly-Gly tripeptide (fMGG) [21]. This model is based on components of an *Escherichia coli* -based reconstituted *in vitro* translation system [16]. We use the chemical reaction network (CRN) formalism to create a detailed mechanistic model using a CRN compiler called BioCRNpyler [24]. The computational model has been developed such that it can synthesize peptides of arbitrary lengths and sequences. Our CRN model initially begins with a minimum of 86 components and creates other intermediate components and reactions based on the sequence of the desired peptide.

BioCRNpyler outputs the model in the standard biological modeling language called the Systems Biology Markup Language (SBML) [34]. The exported SBML files can be simulated with any compatible SBML simulator. We choose to use the BioSCRAPE [29] Python package to simulate the SBML models. The CRN model is converted to an ordinary differential equation by BioSCRAPE and solved using Python `odeint` for desired initial conditions. To convert the CRN to an ODE, each reaction rate is written using the mass-action propensity [35]. We use BioSCRAPE because it supports sensitivity analysis and Bayesian inference tools for SBML models in addition to the model simulations. For each SBML model, we ran local sensitivity analysis to obtain the sensitivity of the measured species with all parameters at all times. Then, we aim to identify the most sensitive parameters for the model using the experimental data. We perform the parameter identification using a Bayesian inference algorithm implemented in BioSCRAPE with emcee Python package [36]. Given the experimental data, we obtain a probability distribution for each identified parameter with Bayesian inference. Model simulations with parameter values sampled from these posterior probability distributions are then plotted against the experimental data to evaluate the quality of the model predictions. These posterior probability distributions also quantify the uncertainty in the data, which is an important advantage of Bayesian inference methods.

Summarizing the computational analysis for the PURE transcription model, we identified four out of eight parameters using the experimental data. The final trained model predicts the transcription from the plasmid’s DNA sequence. In the PURE transcription model, the MGapt signal in the model does not include explicit reactions for aptamer folding or malachite green dye binding. Instead, an MGapt molecule is counted toward the fluorescence signal when RNA_length=64_, the full MGapt transcript length. This assumes that folding and dye binding occur rapidly compared to transcription kinetics, such that transcript completion effectively determines signal generation. The translation kinetic parameters and initial conditions for simulation were taken from the computational model simulating the synthesis of a fMGG, the PURE simulator [21] website (https://sites.google.com/view/puresimulator). The parameter values, initial conditions, and Bayesian inference chains are available in the Supplementary Information.

All calculations and plotting were performed using the standard stack of Python packages – NumPy [37], SciPy [38], Pandas [39], Matplotlib [40], Bokeh [41], and Seaborn [42]. The run time for each simulation depends on the protein length for which the model is created. The simulation time varies from less than a second (for a model with around 200 species and parameters) to a few minutes (for a model with about 3000 species and parameters) on a personal computer running Intel i7-6700 2.6 GHz with 16GB of RAM. The simulation times and Bayesian inference routines can be sped up by around 10x by running the model simulations on a high-performance computing cluster. All data analysis, parameter inference, and data presented in this paper are available on Github at https://github.com/zjuradoq/PURE_CRN_models [43].

### DNA constructs

The original deGFP DNA plasmid, pTXTL-T7p14-mGapt, and pTXTL-T7p14-deGFP were purchased from Arbor Biosciences myTXTL Toolbox 2.0 plasmid collection [44]. The deGFP protein is on a T7 promoter with a UTR1 ribosome binding site (RBS). A MGapt was cloned between the promoter and the RBS site using primers from Integrated DNA Technologies. Forward and reverse primers used can be found in Supplementary Information Table S7.

Subsequently, both plasmids were initially transformed into JM109 cells and cultivated overnight on plates containing carbenicillin resistance at 37 °C. Colonies from each plate were selected and cultured in 4.5 mL LB medium with 4.5 µL of carbenicillin (100 mg*/*µL) overnight. Glycerol stocks of each plasmid were prepared by mixing 500 µL of liquid culture with an equal volume of 50 % glycerol, while the remaining culture was miniprep using the Qiagen Miniprep Kit. Before running DNA in the cell-free reaction, all plasmids underwent an additional PCR purification step utilizing a QiaQuick column (Qiagen) to eliminate excess salt. The purified plasmids were then eluted and stored in nuclease-free water at 4 °C for short-term storage and at −20 °C for long-term storage.

### RNA constructs

Purified RNA was made using the original plasmid of pTXTL-T7p14-mGapt (pT7-MGapt-tT7) and the modified pTXTL-T7p14-deGFP (pT7-MGapt-UTR1-deGFP-tT7) plasmid. Purified plasmids were first amplified and linearized using Q5 Hot Start High-Fidelity DNA Polymerase (NEB); forward and reverse primers are listed in Table S7. Eight reactions of 125 µL, for a total volume of 1 mL were run in a thermocycler. The thermocycler was run for 30 cycles with an elongation time of 15 sec or 45 sec, respectively, and annealing temperature of 65 °C. The completed PCR reactions were then combined into three microcentrifuge tubes of 333 µL to which 33 µL sodium acetate (3 m) and 1 mL Ethanol (100 %) was added. All three tubes were placed at −80 °C for 20 min and then centrifuged at 16 000 xg for 30 min at 4 °C to remove all traces of ethanol. The precipitated DNA was then resuspended in 50 µL of nuclease-free water, combined, and stored at 4 °C.

Using the HiScribe T7 High Yield RNA Synthesis Kit (NEB) 1 µg of the amplified and linearized DNA construct was transcribed in an 80 µL reaction volume and incubated overnight at 37 °C. Before the RNA purification procedure, the transcription reaction underwent DNAse I treatment for 15 min at 37 °C to remove the linear DNA template. Ethanol (100 %) was then added to achieve a final concentration of 35 %, to precipitating the RNA. Next, using PureLink RNA Mini Kit (Invitrogen), RNA was isolated following the protocol for ‘Purification of RNA from liquid samples’, starting from Step 2-Bind RNA. The yield and quality of the RNA samples were analyzed using the NanoDrop2000c, measuring UV absorbance at 260 nm. The purified RNA was flash frozen in liquid nitrogen and stored at −80 °C.

### Dynamic MGapt RNA calibration

Dynamic MGapt RNA calibration was performed to account for the influence of PURE system chemical composition on aptamer fluorescence. As demonstrated by Jurado *et al*. [45], MGapt fluorescence in commercial PURE is significantly affected by DTT concentration and does not directly reflect RNA concentration without correction. The dynamic fluorescence calibration curve for MGapt was generated using purified RNA of MGapt at 0.522 µm and MGapt-UTR1-deGFP at 0.548 µm. A 10 µL reaction was done in triplicate using PURExpress^®^ *In Vitro* Protein Synthesis Kit initially mixed in PCR tubes with 5 % excess. Each reaction contained 10 µm malachite green oxalate, and 8 units of RNase inhibitor. The10 µL samples were loaded to a Nunc 384 well plate and read using a BioTeK H1MF plate reader at 37 °C with 610*/*650 nm (ex/em) and gain 150. The relative fluorescence units (RFU) for each of the RNA units tested are shown in Figure S2.

### Standard deGFP calibration curves

The fluorescence calibration curve for deGFP was generated using purchased purified eGFP from Cell Biolabs (STA-201). Samples were prepared as described in the myTXTL manual [44]. The 1 mg mL^−1^ eGFP (29.0 kDa was estimated to have a concentration of 34.483 µm). The eGFP stock was diluted in series in 1X PBS, and 10 µL of each dilution was pipetted onto the wall of a Nunc 384 well plate, spun down, and then sealed with a plastic film. The plate was allowed to sit for 45 min at room temperature before being read in a BioTek H1MF plate reader at 30 °C and at 485*/*515 nm (ex/em) and gain of 61. Each point on the calibration curve represents the average of 12 points; three replicates were read over 3 minutes at 1-minute intervals to generate 4 points per replicate. The points were all background-subtracted such that the PBS-only samples had zero fluorescence. Points were fit using linear regression and were not forced to go through the origin. Fits for each calibration curve are indicated in the S1.

### PURE reactions and fluorescence measurements

PURE reactions were mixed by following the protocol by PURExpress (E6800), adjusted for a 10 µL reaction, and allowed to run in a 384-well plate (Nunc) at 37 °C. DNA at 5 nm was used, unless otherwise stated, 0.8 units of RNAse inhibitor (NEB), and 10 µm of malachite-green dye was added to each reaction. Fluorescence measurements were read in a Synergy H1 plate reader (Biotek) at 3 min intervals using excitation/emission wavelengths set at 610*/*650 nm (MGapt) at gain 150 and 485*/*515 nm(deGFP) at gain 61. All samples were read in the same plate reader, and for deGFP, relative fluorescence units (RFUs) were converted to nm of protein using a purified eGFP standard by following the protocol in paper [44]. Calibration curves for MGapt and deGFP are depicted in Figure S2 and Figure S1.

## Associated content

### Supporting Information

Supporting Information includes: fluorescence calibration protocols (Section S1), translation model validation and RNA_effective_ derivation (Section S2), Bayesian inference procedures for transcription parameters (Section S3), complete PURE model validation across DNA concentrations (Section S4), and experimental primers (Section S5).

All data analysis, parameter inference, and data presented are available on Github at https://github.com/zjuradoq/PURE_CRN_models.

## Author information

### Author

**Ayush Pandey**–Electrical Engineering and Computer Science, University of California, Merced, California 95343, United States;

**Richard M. Murray**–Division of Engineering and Applied Science, California Institute of Technology, Pasadena, California 91125, United States;

### Author Contributions

Z. J. conceived the project, developed the mathematical model, conducted all experiments and analyzed results, fit the model to the data collected, and wrote the manuscript. A.P. contributed to the development of the model, specifically in the implementation of parameter-fitting algorithms, and to the revision of the manuscript. R. M. M. provided supervision and assisted in editing the manuscript.

### Notes

The authors declare no competing financial interest.

## Acknowledgments

We thank Dr. Samuel Schaffter for the thoughtful review of the manuscript and valuable feedback.

