## Supplementary Material for "Nucleotide-level chemical reaction network modeling enables quantitative prediction of reconstituted cell-free expression system"

February 23, 2026

#### Contents

**1 deGFP Protein and MGapt RNA Calibration** **S3**
1.1 Standard deGFP calibration curves . . . . . S3
1.2 Dynamic MGapt RNA calibration . . . . . S4

**2 Validation of Translation Model with pT7-MGapt-UTR1-deGFP-tT7** **S5**
2.1 Observed relation between RNA and protein . . . . . S5
2.2 Calculation of $RNA_{effective}$ . . . . . S6
2.3 Absolute error of the translation model . . . . . S7

**3 Bayesian Parameter Inference for the Transcription Model** **S7**
3.1 Transcription-only model initial conditions . . . . . S7
3.2 Initial transcription-only model parameters . . . . . S8
3.3 Bayesian inference posterior distributions . . . . . S9
3.4 Transcription with translation model initial conditions . . . . . S10
3.5 Final Parameter Values . . . . . S11

**4 Validation of the PURE Model** **S12**
4.1 Efficiency of PURE . . . . . S12
4.2 Absolute Error of PURE Model . . . . . S13
4.3 Observed relation between DNA, RNA, and protein . . . . . S14
4.4 Calculation of $DNA_{effective}$ . . . . . S15
4.5 Absolute error at different DNA concentrations . . . . . S16

**5 Primers** **S16**

### 1 deGFP Protein and MGapt RNA Calibration

#### 1.1 Standard deGFP calibration curves

The fluorescence calibration curve for deGFP was generated using purchased purified eGFP from Cell Biolabs (STA-201). Samples were prepared as described in the myTXTL manual [1]. The 1 mg mL$^{-1}$ eGFP (29.0 kDa was estimated to have a concentration of 34.483 µM). The eGFP stock was diluted in series in 1X PBS, and 10 µL of each dilution was pipetted onto the wall of a Nunc 384 well plate, spun down, and then sealed with a plastic film. The plate was allowed to sit for 45 min at room temperature before being read in a BioTek H1MF plate reader at 30 °C and at SI485/515nm (ex/em) and gain of 61 (data available on Github at `https://github.com/zjuradoq/PURE_CRN_models` [2]). Each point on the calibration curve represents the average of 12 points; three replicates were read over 3 minutes at 1-minute intervals to generate 4 points per replicate. The points were all background-subtracted such that the PBS-only samples had zero fluorescence. Points were fit using linear regression and were not forced to go through the origin. Fits for each calibration curve are indicated in the Figure S1.

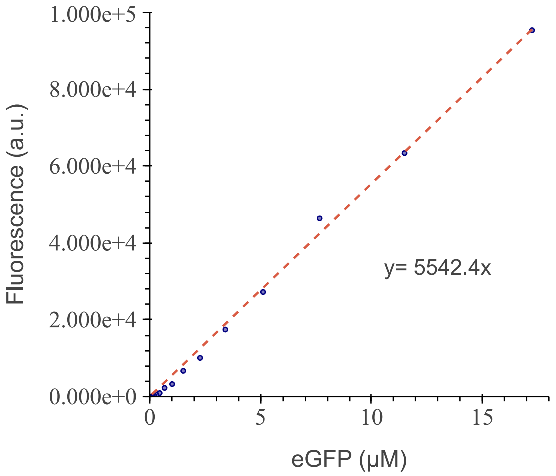


**Figure S1: Standard deGFP calibration curve.** The fluorescence calibration curve for deGFP is used to convert RFU to µM.

#### 1.2 Dynamic MGapt RNA calibration

MGapt fluorescence calibration accounts for DTT concentration and chemical composition effects on aptamer signal in commercial PURE systems Jurado *et al.* [3]. The dynamic fluorescence calibration curve for MGapt was generated using purified RNA of MGapt at 0.522 µM and MGapt-UTR1-deGFP at 0.548 µM in a 10 µL PURExpress reaction done in three technical replicates. The PURE reaction with the purified RNA containing the MGapt was loaded to a Nunc 384 well plate and was read using a BioTeK H1MF plate reader at 37 °C and at SI610/650nm (ex/em) and gain 150. The relative fluorescence units (RFU) for each of the RNA units tested are shown in Figure S2a (MGapt) and Figure S2b (MGapt-UTR1-deGFP). Subsequently, dynamic calibration curves for RNA of MGapt (Figure S2c) and MGapt-UTR1-deGFP (Figure S2d) were obtained by dividing RFU measurements in Figure S2a and Figure S2b by their respective RNA concentrations, then smoothed. The dynamic fluorescence calibration curve for MGapt is specific to the PURE reaction and the RNA produced.

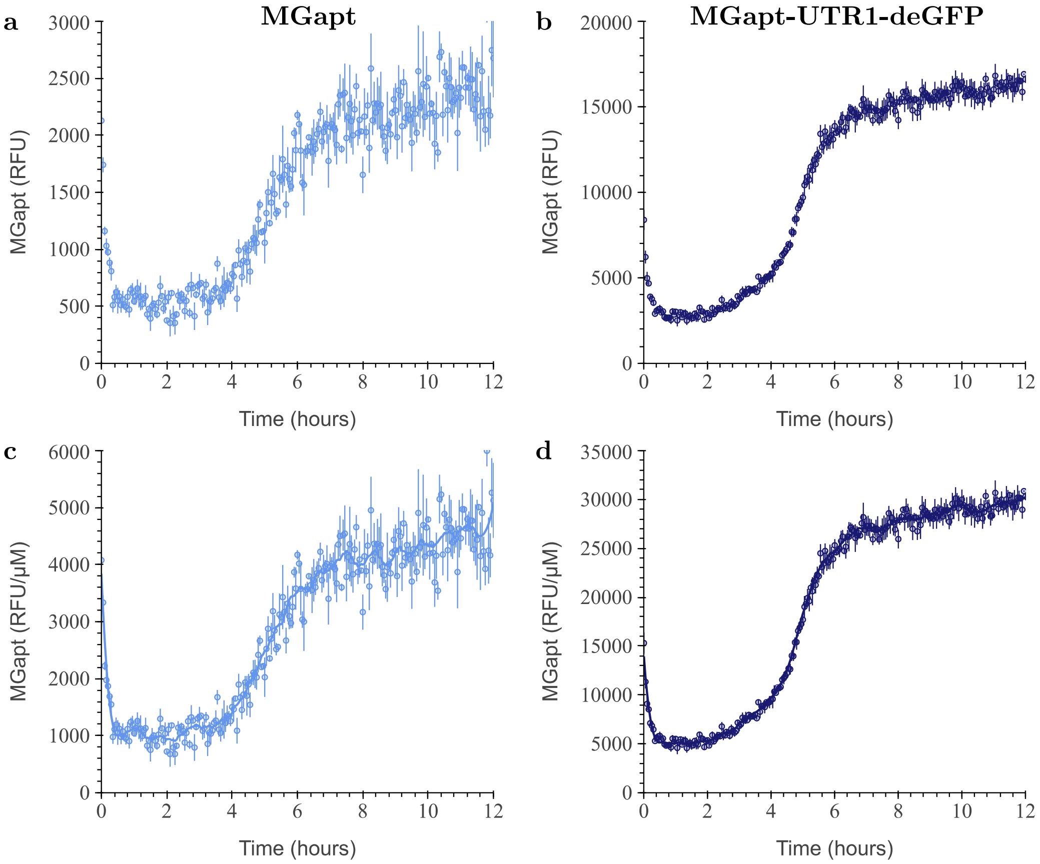


**Figure S2: Dynamic MGapt calibration curve.** Measured RFU of RNA of (**a**) MGapt at 0.522 µM and (**b**) MGapt-UTR1-deGFP at 0.548 µM done in triplicate (circles with error bars). Respective dynamic calibration curves for (**c**) MGapt and (**d**) MGapt-UTR1-deGFP RNA, calculated by dividing RFU measurement in (**a**) and (**b**) by respective RNA concentration, then smoothing.

### 2 Validation of Translation Model with pT7-MGapt-UTR1-deGFP-tT7

#### 2.1 Observed relation between RNA and protein

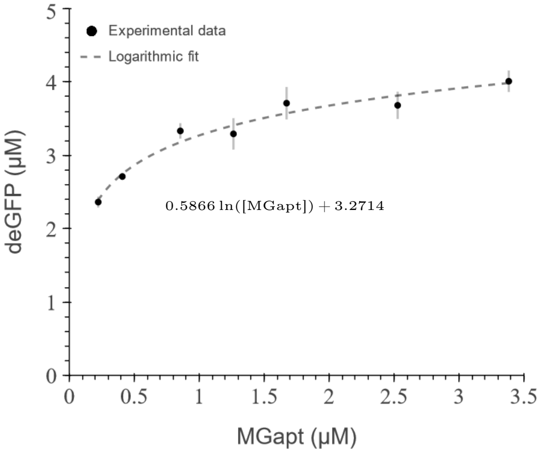


**Figure S3: Relationship between deGFP production and initial RNA concentrations.** Translation was initialized by the addition of RNA to achieve final concentrations: 0.22 μM, 0.41 μM, 0.86 μM, 1.23 μM, 1.67 μM, 2.53 μM, and 3.38 μM. The corresponding average deGFP production (black circles with associated error bars) was fit using a log fit due to RNA's relationship with deGFP production. The fitted line overlays the data (dashed black line) with the equation displayed.

The relationship between deGFP production and initial RNA is given by the equation,

$$[\text{deGFP}] = 0.5866 \ln([\text{MGapt}]) + 3.2714. \tag{3.9}$$

#### 2.2 Calculation of $RNA_{effective}$

Effective RNA ($RNA_{effective}$) is given by the equation,

$$RNA_{effective} = k\,(RNA) \cdot RNA. \qquad (3.10)$$

To calculate the multiplication factor $k\,(RNA)$ in equation (3.10) we identified the effective RNA ($RNA_{effective}$) that produces the corresponding deGFP for RNA concentrations at 0.22 µM and 3.38 µM (see Figure S4a). Next, we compute $k\,(RNA)$ at 0.22 µM and 3.38 µM. Finally, shown in Figure S4b, we plot the computed $k\,(RNA)$ against MGapt concentration and imposing a power trendline, we fit the points giving $k\,(RNA){=}0.1703\,RNA^{-0.801}$. We imposed a power law relationship to capture the observed nonlinear decline in translation efficiency with increasing RNA concentration. This functional form empirically describes the saturation behavior where protein production per RNA molecule decreases as total RNA increases, consistent with resource competition and machinery loading effects in PURE.

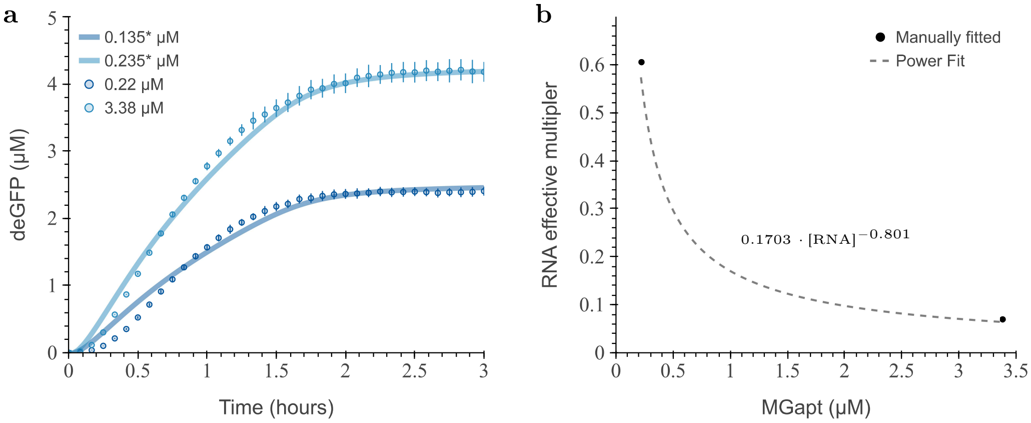


**Figure S4: Effective RNA required for the model to achieve corresponding deGFP production.** (**a**) Results from manually tuning the initial RNA concentrations to match the deGFP production from the two ends of the RNA concentration experimentally tested: 0.22 µM, and 3.38 µM (circles with error bars). The model results with manually tuned RNA are shown by solid lines with colors corresponding to the experimental data. (**b**) The proposed RNA multiplier function to calculate the effective RNA concentrations to fit experimental deGFP production with the translation model.

#### 2.3 Absolute error of the translation model

**Table S1:** Absolute error of the translation model compared to experimental results. The table lists the initial RNA conditions not used in determining the effective RNA multiplier function. The deGFP expressed reflects the concentrations at 2 h for both the simulation and experiment.

| $RNA_{added}$ | $RNA_{effective}$ | $deGFP_{experimental}$ | $deGFP_{simulated}$ | Absolute Error |
|---|---|---|---|---|
| 0.409 µM | 0.153 µM | 2.71 µM | 2.65 µM | 2.21 % |
| 0.855 µM | 0.178 µM | 3.33 µM | 3.08 µM | 7.51 % |
| 1.264 µM | 0.192 µM | 3.29 µM | 3.31 µM | 0.61 % |
| 1.67 µM | 0.204 µM | 3.71 µM | 3.50 µM | 5.66 % |
| 2.528 µM | 0.221 µM | 3.68 µM | 3.78 µM | 2.72 % |

### 3 Bayesian Parameter Inference for the Transcription Model

#### 3.1 Transcription-only model initial conditions

To account for the energy usage of translation-related reactions separate from protein production, we modified the initial conditions used in the Bayesian inference pipeline for the transcription model of DNA construct pT7-MGapt-tT7.

**Table S2:** Initial conditions used in the Bayesian inference pipeline for the transcription-only model.

| Species | Value | Unit |
|---|---|---|
| ATP | 2206 | µM |
| GTP | 590 | µM |
| CTP | 1250 | µM |
| UTP | 1250 | µM |
| DNA | 5 | nM |
| T7 RNAP | 1 | µM |

#### 3.2 Initial transcription-only model parameters

The initial parameter values for the transcription model are given in Table S3. The initial chemical reaction rates of the transcription model were based initially on the TX-TL model by Tuza *et al.* [4].

**Table S3:** Initial transcription model parameters for PURE cell-free expression.

| Parameter | Description | Value | Unit |
|---|---|---|---|
| $k_1$ | Binding of RNAP and GTP to the DNA | 6.10 | $\mu M^{-2}\,s^{-1}$ |
| $k_2$ | Rate of formation of the RNAP bound GDP and phosphate complex on the DNA from RNAP bound GTP complex | 2.95 | $s^{-1}$ |
| $k_3$ | Unbinding of GDP and Phosphate from the RNAP and DNA complex | 7.82 | $s^{-1}$ |
| $k_{start}$ | Start of the initiation of the RNA transcript, ($RNA_0$) from the RNAP and DNA complex | 5.24 | $s^{-1}$ |
| $k_{NTP_{bound}}$ | Binding rate of NTP to the RNAP bound DNA, complex with initiated RNA transcript | 1.47 | $\mu M^{-1}\,s^{-1}$ |
| $k_{NTP_{add}}$ | Rate of elongation of the transcript | 23.59 | $s^{-1}$ |
| $k_{NTP_{dis}}$ | Unbinding rate of NMP and PPi from the open complex | 985.89 | $s^{-1}$ |
| $k_{term}$ | Termination rate | 32.38 | $s^{-1}$ |

#### 3.3 Bayesian inference posterior distributions

The initial posterior distributions of parameters, shown in Figure S5, were obtained using the Bayesian inference tools in Bioscrape on all the reaction rates. Subsequently, the model was retrained using a narrower standard deviation around the results from the initial inference, shown in Figure 4.

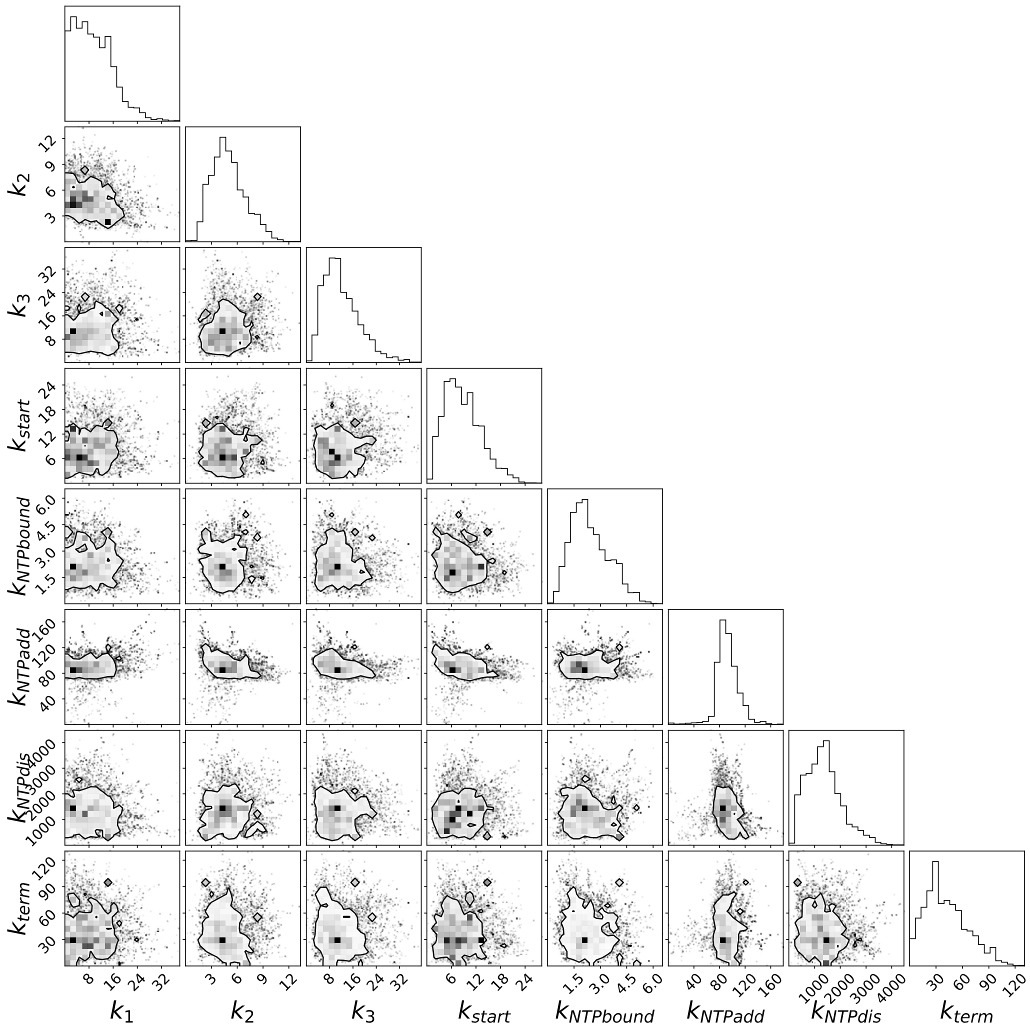


**Figure S5: Initial coarse Bayesian inference posterior distributions.** The initial posterior distributions of parameters on all eight reaction rates. The corner plot depicts the covariance of the eight parameters, with the contour showing the 75 % probability region for the parameter values having limited information on accurate parameter values.

#### 3.4 Transcription with translation model initial conditions

The initial conditions for the combined transcription and translation model are given in Table S4; based on Version 7 PURE concentration published in Table S1 by Kazuta *et al.* [5]

**Table S4:** Translation initial condition for the 36 proteins of PURE cell-free reaction model.

| Species | Value | Unit |
|---|---|---|
| ATP | 3750 | µM |
| GTP | 2500 | µM |
| CTP | 1250 | µM |
| UTP | 1250 | µM |
| DNA | 5 | nM |
| CP | 10 | mM |
| FD | 126.8498943 | µM |
| T7 RNAP | 1 | µM |
| CK | 10 | µM |
| EFG | 4.3 | µM |
| EFTs | 13 | µM |
| EFTu | 80 | µM |
| IF1 | 99 | µM |
| IF2 | 4.1 | µM |
| IF3 | 4.9 | µM |
| MK | 5.6 | µM |
| MTF | 2.4 | µM |
| NDK | 1.8 | µM |
| PPiase | 0.16 | µM |
| RF1 | 0.2 | µM |
| RF2 | 0.2 | µM |
| RF3 | 0.7 | µM |
| RRF | 16 | µM |

| Species | Value | Unit |
|---|---|---|
| AAs | 300 | µM |
| AlaRS | 3 | µM |
| ArgRS | 0.12 | µM |
| AsnRS | 1.7 | µM |
| AspRS | 0.49 | µM |
| CysRS | 0.1 | µM |
| GlnRS | 0.24 | µM |
| GlyRS | 0.35 | µM |
| GluRS | 0.9 | µM |
| HisRS | 0.34 | µM |
| IleRS | 1.5 | µM |
| LeuRS | 0.16 | µM |
| LysRS | 0.46 | µM |
| MetRS | 0.44 | µM |
| PheRS | 0.54 | µM |
| ProRS | 0.67 | µM |
| SerRS | 0.16 | µM |
| ThrRS | 0.34 | µM |
| TrpRS | 0.11 | µM |
| TyrRS | 0.03 | µM |
| ValRS | 0.07 | µM |
| RS70S | 3 | µM |

#### 3.5 Final Parameter Values

The parameter values for the transcription model are given in Table S5. The initial chemical reaction rates of the transcription model were based on the TX-TL model by Tuza *et al.* [4].

**Table S5:** Final transcription model parameters for PURE cell-free extract.

| Parameter | Description | Value | Unit |
|---|---|---|---|
| $k_1$ | Binding of RNAP and GTP to the DNA | 9.41 | $\mu M^{-2} s^{-1}$ |
| $k_2$ | Rate of formation of the RNAP bound GDP and phosphate complex on the DNA from RNAP bound GTP complex | 5.06 | $s^{-1}$ |
| $k_3$ | Unbinding of GDP and Phosphate from the RNAP and DNA complex | 10.88 | $s^{-1}$ |
| $k_{start}$ | Start of the initiation of the RNA transcript, ($RNA_0$) from the RNAP and DNA complex | 7.64 | $s^{-1}$ |
| $k_{NTP_{bound}}$ | Binding rate of NTP to the RNAP bound DNA, complex with initiated RNA transcript | 2.68 | $\mu M^{-1} s^{-1}$ |
| $k_{NTP_{add}}$ | Rate of elongation of the transcript | 102.17 | $s^{-1}$ |
| $k_{NTP_{dis}}$ | Unbinding rate of NMP and PPi from the open complex | 1306.62 | $s^{-1}$ |
| $k_{term}$ | Termination rate | 45.99 | $s^{-1}$ |

The model's number of parameters depends on the transcript sequence. For example, the transcription model for malachite green aptamer has 276 parameters.

### 4 Validation of the PURE Model

#### 4.1 Efficiency of PURE

The synthesis of incomplete peptides from the DNA construct pT7-UTR1-deGFP-tT7 can be observed using radiolabeled [$^{35}$S]-methionine [6] in two cell-free protein synthesis systems. The synthesis of full-length deGFP is approximately 28 kDa, and thus should measure between 26 kDa and 34 kDa. The completed translation of deGFP is denoted by the dark band observed in both BL21 (DE3) *E. coli* cell-lysate (Figure S6a) and NEB PURExpress (Figure S6b).

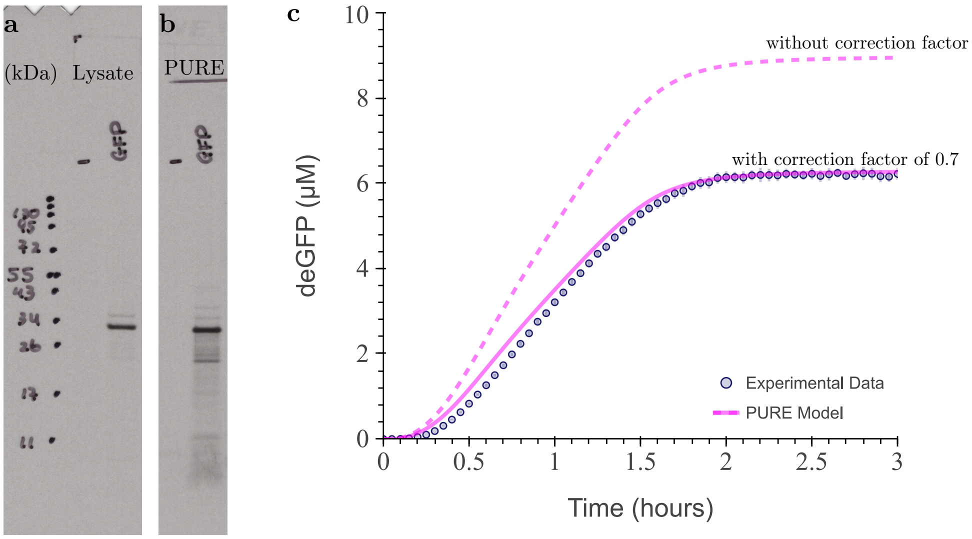


**Figure S6: [$^{35}$S]-methionine labeling of lysate-based and PURE cell-free protein synthesis systems.** Newly synthesized proteins from the DNA construct pT7-UTR1-deGFP-tT7 using BL21 (DE3) *E. coli* cell-lysate (**a**) and NEB PURExpress (**b**) using radiolabeled [$^{35}$S]-methionine. (**c**) Modeled deGFP expression in the combined models (magenta line) with correction factor of 0.7 (solid line) and without (dashed line) overlaying with experimental data, three replicates (blue circles and blue error bars). Figures (a) and (b) are courtesy of Masami Hazu, a PhD candidate in Prof. Voorhees' lab, who collaborated with us to conduct the experiment and contributed to the final results.

#### 4.2 Absolute Error of PURE Model

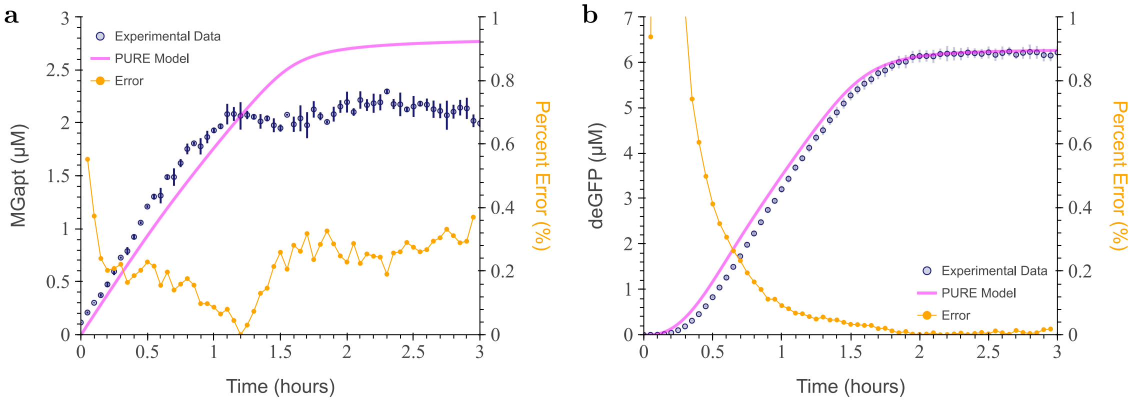


**Figure S7: Absolute error of combined PURE model compared to experimental results.** (**a**) Modeled RNA production in the combined model (magenta line) overlayed with experimental data, three replicates (blue circles and blue error bars), and the absolute error (orange circles) on a secondary axis. (**b**) Modeled deGFP expression in the combined model (magenta line) overlayed with experimental data, three replicates (blue circles and blue error bars), and the absolute error (orange circles) on a secondary axis.

#### 4.3 Observed relation between DNA, RNA, and protein

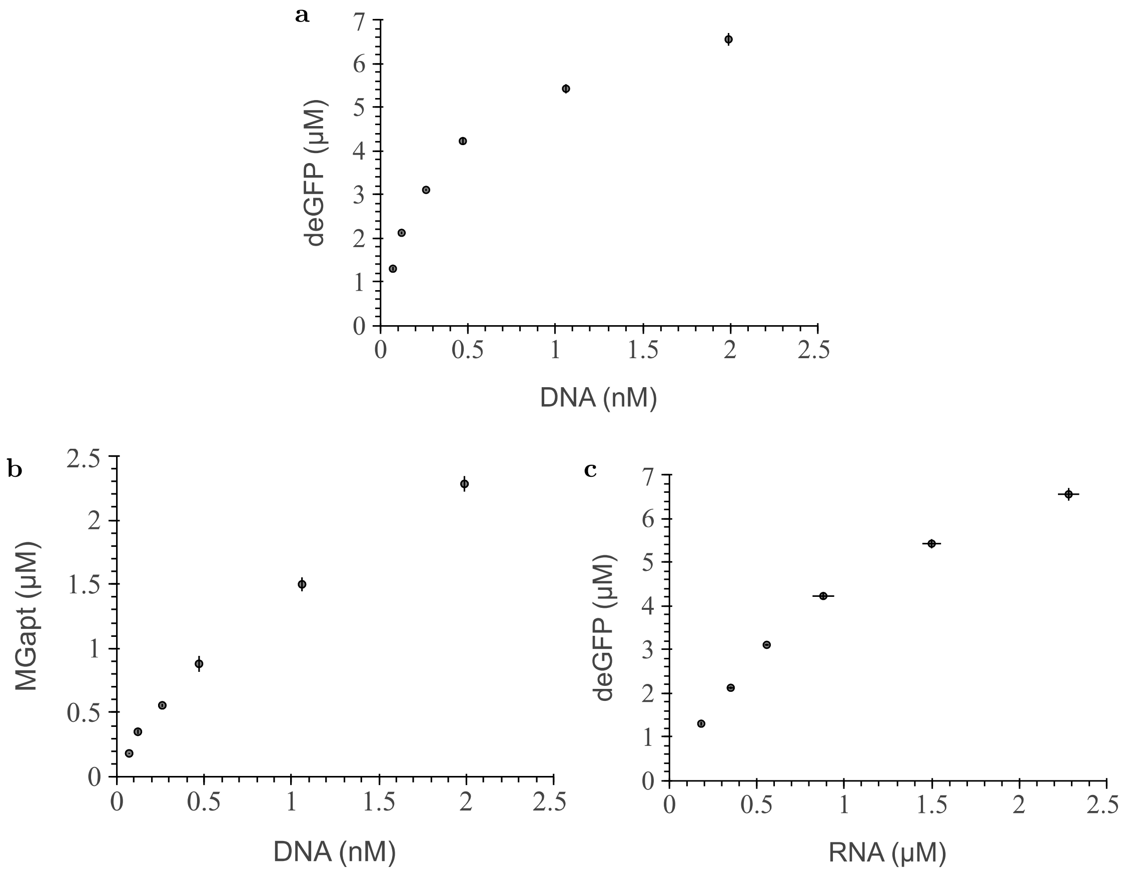


**Figure S8: Relationship between deGFP and RNA production at varying initial DNA concentrations.** Expression from the plasmids of pT7-MGapt-UTR1-deGFP-tT7 at DNA the following concentrations: 0.07 nM, 0.12 nM, 0.26 nM, 0.47 nM, 1.06 nM, and 1.99 nM. (**a**) The initial DNA concentration versus average overall deGFP production at 2 h is depicted, followed by a breakdown into (**b**) RNA production relative to DNA and (**c**) deGFP production based on RNA production. The corresponding experimental data for each subplot is plotted in black circles with associated error bars.

#### 4.4 Calculation of $\mathrm{DNA}_{\mathrm{effective}}$

The effective DNA ($\mathrm{DNA}_{\mathrm{effective}}$) given by the equation,

$$\mathrm{DNA}_{\mathrm{effective}} = k\,(\mathrm{DNA}) \cdot \mathrm{DNA}. \tag{3.12}$$

To calculate the multiplication factor $k\,(\mathrm{DNA})$ in equation (3.12) we identified the effective DNA ($\mathrm{DNA}_{\mathrm{effective}}$) that produces the corresponding RNA for DNA concentrations at 0 nM, 0.12 nM, 0.47 nM, and 1.99 nM. Next, we plot the $\mathrm{DNA}_{\mathrm{effective}}$ against the experimental DNA concentration. We used a linear regression to fit the points when DNA was below 0.5 nM and a logarithmic trendline to fit the points above 0.5 nM, as shown in Figure S9.

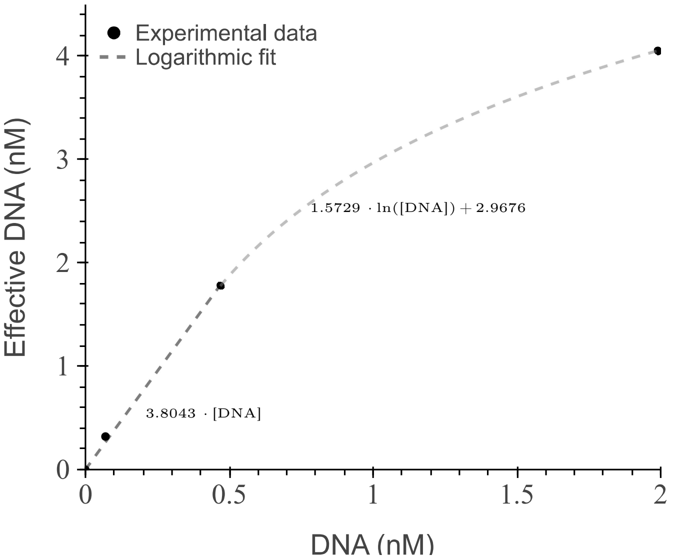


**Figure S9: Effective DNA required for the model to achieve corresponding RNA production.** The proposed piece-wise DNA function calculates effective DNA concentrations needed to fit experimental RNA production with the combined transcription and translation model.

##### 4.5 Absolute error at different DNA concentrations

The error between simulation and experiment at 2 h, shown in Figure 7, is summarized below.

**Table S6:** Absolute error of MGapt and deGFP production of the PURE model over multiple DNA concentrations. The table includes the tested initial DNA conditions, corresponding effective DNA, MGapt synthesized, and protein production results from the model and experimental data. The MGapt and deGFP expressed reflect the concentrations at 2 h for both the simulation and experiment.

| | | | | | | |
|---|---|---|---|---|---|---|
| $DNA_{added}$ (nM) | 1.99 | 1.06 | 0.47 | 0.26 | 0.12 | 0.07 |
| $DNA_{effective}$ (nM) | 4.05 | 3.06 | 1.79 | 0.99 | 0.46 | 0.27 |
| $MGapt_{exp.}$ (µM) | 2.28 | 1.5 | 0.88 | 0.56 | 0.35 | 0.18 |
| $MGapt_{model}$ (µM) | 2.26 | 1.77 | 1.09 | 0.63 | 0.30 | 0.18 |
| $MGapt_{error}$ (%) | 0.88 | 18.0 | 23.9 | 12.5 | 14.3 | 0.0 |
| $deGFP_{exp.}$ (nM) | 6.55 | 5.42 | 4.23 | 3.11 | 2.13 | 1.31 |
| $deGFP_{model}$ (µM) | 5.77 | 5.20 | 3.91 | 2.55 | 1.31 | 0.79 |
| $deGFP_{error}$ (%) | 11.9 | 4.06 | 7.57 | 18.0 | 38.5 | 39.7 |

### 5 Primers

Primers used to clone MGapt (GGGATCCCGACTGGCGAGAGCCAGGTAACGAATGGATC) into DNA plasmid, pTXTL-T7p14-deGFP, obtained initially from myTXTL [1] and to linearize DNA for RNA purifications.

**Table S7:** List of primers used to make constructs.

| Name | Sequence | Purpose |
|---|---|---|
| pT7_MGapt_FOR | GAGCCAGGTAACGAATGGATCCAAT**A ATTTTGTTTAACTTTAAGAAGGA GATATACCATG** | Cloning in MGapt to pTXTL-T7p14-deGFP |
| pT7_MGapt_REV | ATTGGATCCATTCGTTACCTGGCTCTC GCCAGTCGGGATCCC**TCTAGAGGGA AACCGTTG** | Cloning in MGapt to pTXTL-T7p14-deGFP |
| pPCR_MGapt_FOR | **GTGATGTCGGCGATATAGGC** | Linearize pTXTL-T7p14-mGapt |
| pPCR_MGapt_REV | **CACTATCGACTACGCGATCATG** | Linearize pTXTL-T7p14-mGapt |
| pPCR_MGapt-UTR1 -deGFP_FOR | **GCGTAGAGGATCGAGATCTCGAT C** | Linearize modified pTXTL-T7p14-deGFP |
| pPCR_MGapt-UTR1 -deGFP_REV | **CTATCGACTACGCGATCATGGC** | Linearize modified pTXTL-T7p14-deGFP |

The bold text identifies the binding region of the plasmid.

### References

[1] Arbor Biosciences. *myTXTL T7 Expression Kit*. Michigan, United States of America, 2019.

[2] Zoila Jurado. Source code for PURE CRN models. `https://github.com/zjuradoq/PURE_CRN_models`, 2024.

[3] Zoila Jurado and Richard M. Murray. Impact of chemical dynamics of commercial PURE systems on malachite green aptamer fluorescence. *ACS Synthetic Biology*, 2024.

[4] Zoltán A. Tuza, Dan Siegal-Gaskins, Jongmin Kim, and Gábor Szederkényi. Analysis-based parameter estimation of an *in vitro* transcription-translation system. In *Eur. Control Conf. 2015*, page 1554–1560, 2015.

[5] Yasuaki Kazuta, Tomoaki Matsuura, Norikazu Ichihashi, and Tetsuya Yomo. Synthesis of milligram quantities of proteins using a reconstituted in vitro protein synthesis system. *Journal of Bioscience and Bioengineering*, 118(5):554–557, 2014.

[6] Lin-Chi Chen and Arturo Casadevall. Labeling of proteins with [$^{35}$s] methionine and/or [$^{35}$s] cysteine in the absence of cells. *Analytical Biochemistry*, 269(1):179–188, 1999.
